# Time-lapse single-cell transcriptomics reveals modulation of histone H3 for dormancy breaking

**DOI:** 10.1101/720680

**Authors:** Hayato Tsuyuzaki, Masahito Hosokawa, Koji Arikawa, Takuya Yoda, Naoyuki Okada, Haruko Takeyama, Masamitsu Sato

## Abstract

How quiescent cells break dormancy is a key issue in eukaryotic cells including cancer. Fungal spores, for example, remain quiescent for long periods until nourished, although the mechanisms by which dormancy is broken remain enigmatic. Transcriptome analysis could provide a clue, but methods to synchronously germinate large numbers of spores are lacking, and thus it remains a challenge to analyse gene expression upon germination. Hence, we developed methods to assemble transcriptomes from individual, asynchronous spore cells of fission yeast undergoing germination to assess transcriptomic changes over time. The virtual time-lapse analyses highlighted one of three copies of histone H3 genes for which transcription fluctuates during the initial stage of germination. Disruption of this temporal fluctuation caused defects in spore germination despite no visible defects in other stages of the life cycle. We conclude that modulation of histone H3 expression is a crucial ‘wake-up’ trigger at dormancy breaking.

## Main Text

### Results

#### Establishing single-cell transcriptomes

Eukaryotic cells of fungi, plants and animals often exist in a non-dividing state called quiescence or dormancy. However, cells may begin to proliferate upon a change in environmental conditions such as nutrient availability or signaling from neighbouring cells. The fission yeast *Schizosaccharomyces pombe* undergoes meiosis and gametogenesis under nutrition starvation to generate round-shaped spores that remain dormant until nutrients become available (**Fig. 1a**). Nutrition refeeding gradually breaks spore dormancy, and germinating spores undergo morphological changes such as cell-wall expansion accompanied by cytoskeletal rearrangements to form a germ projection (**Fig. 1a**) ^1^. The cell cycle is then activated.

**Fig. 1.**
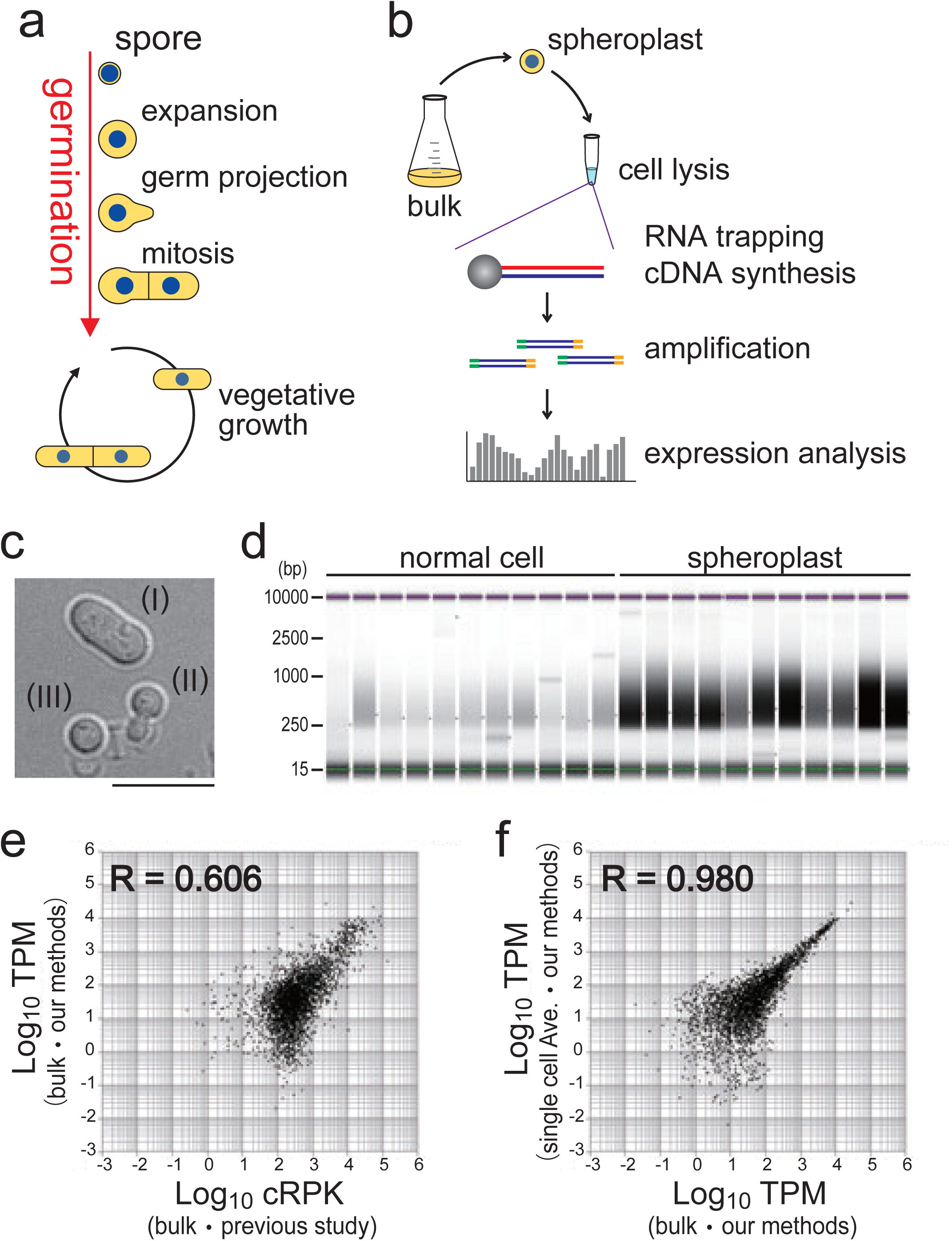
Validation of Single-cell Transcriptome Methods. **(a)** Germination processes and entry into vegetative growth of *S. pombe* cells. **(b)** Overview of the bead-based scRNA-seq developed in this study. **(c)** Differential interference-contrast images of WT cells after treatment with lysing enzyme: (I) cells not lysed, (II) cells being lysed, and (III) cells that were substantially lysed. Scale bar, 10 µm. **(d)** Assessment of scRNA-seq library from 22 single cells for which spheroplasts were prepared or not prepared (Normal cell), as assessed with electrophoresis. Each lane corresponds to a sample from a single cell. Size (bp) is shown to the left. **(e)** Comparison of *S. pombe* transcriptomes created from bulk cells in this study and in a previous study ^5^. **(f)** Comparison of single cell–based transcriptomes (average of 11 cells) and transcriptomes created from bulk cells in this study. TPM: transcripts per million mapped reads, cRPK: reads per kilobase of length corrected for mappability.

It is likely that these dormancy-breaking events are brought about by alterations in gene expression upon feeding, although the genes involved have not been fully elucidated. Therefore, defining the temporal changes in gene-expression profiles would provide direct evidence concerning the mechanism of germination. The creation of transcriptional profiles for cells at germination, requires that total cellular RNA be harvested from a large number of cells undergoing germination ^2^. It appears, however, that individual cells tend to break dormancy at a distinct time, and no methods have been established to synchronously induce spore germination for bulk spore cells.

To overcome this technical limitation, we carried out single-cell transcriptome profiling (single-cell RNA sequencing; scRNA-seq) to create transcriptional profiles from single cells at the state of our interest (such as a germinating spore); this approach bypasses the need for synchronised cultures. The overall strategy is presented in **Fig. 1b**. As there are few examples of single cell–based genomic analyses in yeast ^3,4^, we first determined if we could faithfully reproduce the transcriptomes of vegetative cells in comparison with transcriptomes derived from bulk cultures using standard methods ^5^. First, we utilised spheroplasting of cells prepared in bulk culture to effect severe damage of the cell wall, as indicated by a change to a round cell shape (**Fig. 1c**). Each single cell was picked up and then chemically lysed to extract poly(A)+ RNAs, which were then captured with nanoparticle beads (see **Fig. 1b**), on which reverse transcription and subsequent procedures were performed so that double-stranded cDNAs could be prepared. We particularly applied the “bead-seq” method ^6^ to minimise biased amplification of cDNAs and thereby reduce the background noise during data analysis. For a quality check, cDNA library preparation was confirmed through electrophoresis (**Fig. 1d**). The cDNA library pool was then deep sequenced and mapped on the reference genome to assemble a transcriptome (scRNA-seq).

To evaluate whether our methods generated high-quality transcriptomes, we compared our single cell–based scRNA-seq transcriptome with a published profile created from bulk cells using conventional methods ^5^. When the data for each of ∼7000 *S. pombe* annotated genes was plotted versus the expression level measured in those two distinct transcriptomes, the correlation coefficient was R=0.606, indicating that, on average, gene expression was detected at similar levels in the two transcriptomes, although there were certain genes (**Fig. 1e**). This may be attributable to ‘individuality’ in the expression of genes in individual cells or to differences in protocols. A comparison of profiles created from individual cells and bulk (∼10) cells using our same methods, revealed only subtle differences (R = 0.980, **Fig. 1f**), indicating that any difference with published profiles (**Fig. 1e**) could be attributed to technical differences and that our scRNA-seq protocol for single-cell transcriptome assembly yielded results comparable with those from studies of bulk-cells (**Fig.1f**).

#### Expanding the scope of the protocol to single spores

Our protocol was then applied to create single-spore transcriptomes, but a sufficient amount of RNA could not be extracted from a single spore cell, probably because the hard spore wall hampered cell lysis (see below). To circumvent this issue, we engineered the Pnmt41-*bgs2* strain to reduce expression of β-glucan synthase, which is required specifically for spore-wall synthesis but not for cell-wall integrity during vegetative growth ^7^. In standard medium, the Pnmt41-*bgs2* strain produced spores with a fragile spore wall (**Fig. 2a**), and indeed the spores burst when suspended in water (**Fig. 2a**). Thus, spore lysis could be achieved without using lysing enzymes, which is a great technical advantage for enabling single-spore omics.

**Fig. 2.**
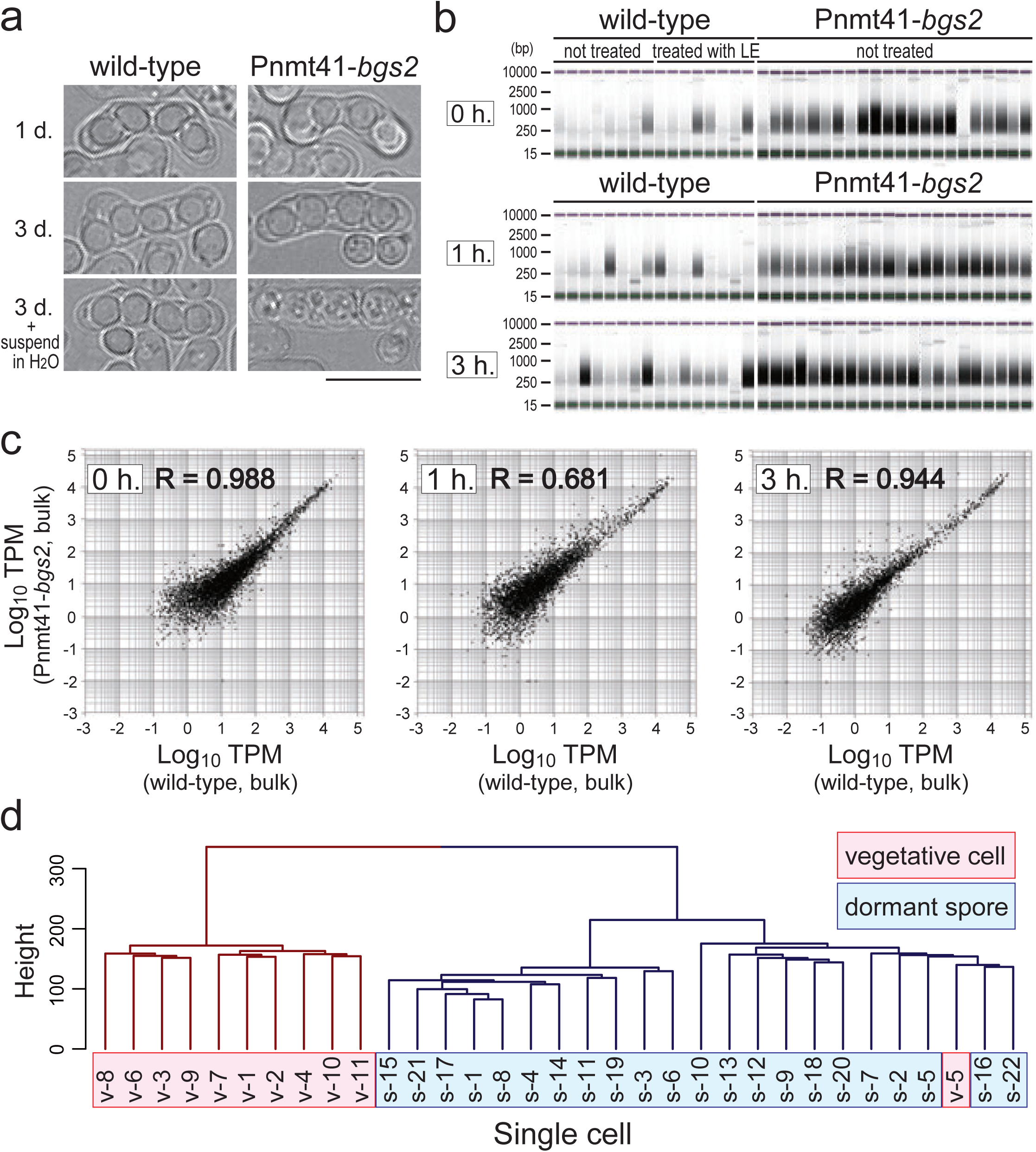
scRNA-seq-based transcriptome assembly from a single spore cell. Strain Pnmt41-*bgs2* was used to efficiently prepare cDNAs from dormant or germinating spores. **(a)** Strain Pnmt41-*bgs2* undergoes meiosis but produces fragile spores. WT (*bgs2*^+^) spores could tolerate to water, whereas Pnmt41-*bgs2* spores burst. **(b**,**c)** Spores of the indicated genotypes were picked at 0, 1 and 3 h after nutrition feeding. **(b)** cDNAs were not produced in sufficient amounts from WT cells irrespective of whether they were treated with the lysing enzyme (left), whereas cDNA could be produced in sufficient amounts from Pnmt41-*bgs2* spores even without lysing enzyme treatment (right). **(c)** Global gene expression measured in strain Pnmt41-*bgs2* did not differ significantly from that of the WT cells. **(d)** Transcription profiles of single vegetative cells (v-1–v-11) and unfed dormant spores (s-1–s-22) were classified into distinct groups through an informatic operation. Relative height reflects the similarity between transcriptomes.

Wild-type and Pnmt41-*bgs2* spores were fed with nutrients to induce germination, and single spores were picked at 0, 1 and 3 h and then burst in water. RNA-seq library was more efficiently amplified from single cells of Pnmt41-*bgs2* spores than from wild-type (WT) spores (**Fig. 2b**). In addition, the reduction in Bgs2 expression in Pnmt41-*bgs2* did not affect the overall transcription profile in dormant or germinating spores (**Fig. 2c** and **Supplementary Fig. 1**). Thus, the improved method was validated for profiling gene expression in a single spore.

Having established experimental procedures, the transcriptomes of single spores were compared to those of single vegetative cells (**Fig. 2d**). A total of 33 single-cell transcriptomes from vegetative cells and unfed dormant spores were sorted out into two major groups. One group exclusively comprised vegetative profiles, and the other comprises profiles mostly from dormant spores (**Fig. 2d**), verifying that our single-cell methodology is sensitive enough to detect changes in transcription in cells exposed to different nutrient conditions.

#### Estimating temporal changes in gene expression

To specifically focus on genes for which expression varied during the initial stage of germination, it was necessary to determine which transcriptome was derived from an individual spore at that stage. This was impossible to deduce based on the appearance of spores when they were picked, as spores at the initial stage were morphologically similar. For that purpose, we collected 64 spores at 0, 1 and 3 h after induction of germination by nutrition refeeding (20 ∼ 22 spores from each timepoint) and assembled transcriptomes for each spore. To determine which transcriptomes corresponded to spores that were at the initial stage of germination, we implemented the “monocle” operation (see below) to compare differences in all 64 transcriptomes, thereby deducing the temporal shift in transcriptomes from one to another ^8–10^. Variation between any two transcriptomes implies a temporal shift from one to another, and therefore any small variation could be derived from two successive transcriptomes over time. Repeating the monocle operation thus enables the clustering of all transcriptomes, from which changes in transcription profiles can be estimated over time, facilitating the temporal alignment of profiles to produce a virtual timeline.

**Fig. 3a** illustrates how the monocle operates: we applied the monocle operation to an imaginary transcriptome comprising expression data for three genes, namely ‘a’, ‘b’, and ‘c’, in terms of transcripts per million reads (TPM) (**Fig. 3a-I**). In this case, each transcriptome can be plotted in three-dimensional space depending on the expression of each gene. The monocle operation projects the three-dimensional location of each transcriptome onto a two-dimensional plane (**Fig. 3a-II,a-III**). As spatial distance on the two-dimensional plane reflects temporal distance, the monocle operation estimates the timeline along with spatial proximity data (**Fig. 3a-IV**).

**Fig. 3.**
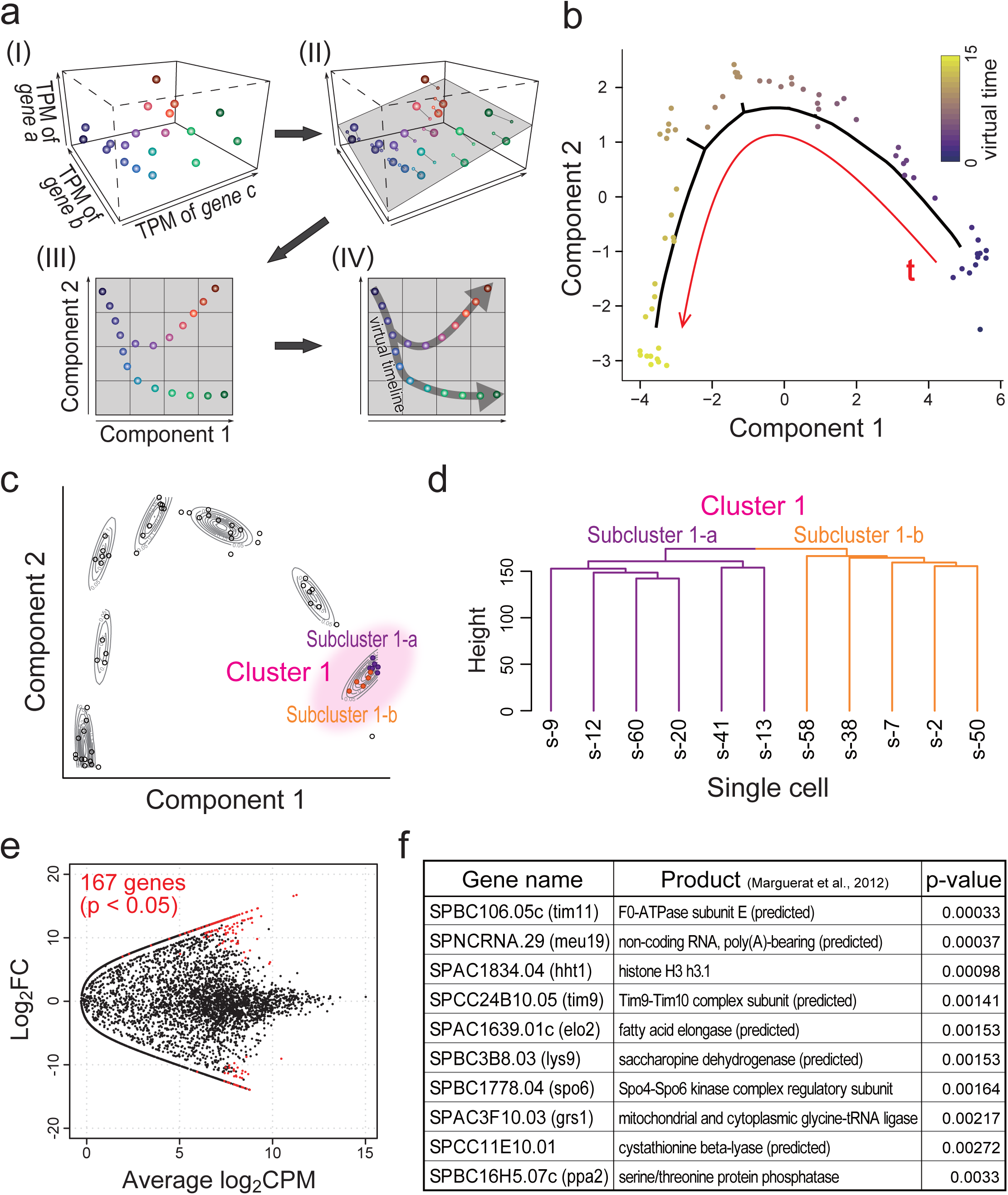
Transcriptomes could be aligned along a virtual timeline. Bioinformatics approaches were applied to estimate temporal changes in gene expression profiles during germination. **(a)** Schematics illustrating the idea of the monocle-based similarity judgment. (I) Assuming that a cell contains three genes (a–c). In this example, transcriptomes can be plotted in three-dimensional space (dots in the graph) depending on the extent to which these genes were expressed. (II) Those data points can be projected onto a single plane (grey) so that the number of dimensions can be reduced from 3 to 2, as depicted in (III). In the two-dimensional plane, the Euclidean distance between data points indicates the similarity between two transcriptomes, and this similarity is used to estimate the time-flow the virtual timeline through a minimum spanning tree (IV). **(b)** Two-dimensional transcriptome plots, which were compressed from ∼7000 dimensions using repetitive monocle operations. We used 64 samples created from single spores before and after nutrition feeding (0, 1, and 3 h). The red curve indicates the virtual timeline, and transcriptomes were aligned in the time course indicated by the heat map. **(c)** Plots in **(b)** were classified into seven clusters along the virtual timeline. Cluster 1 corresponds to the very initial stage of germination estimated from the timeline, and plots therein could be further divided into two subclusters (1-a, purple; and 1-b, orange). **(d)** Transcriptomes in two subclusters were divided into distinct groups through an informatic operation. **(e)** The expression of ∼7000 annotated genes (dots) was compared between subcluster 1-a and 1-An exact test (p < 0.05) revealed 167 genes for which expression varied significantly between these subclusters (shown in red). **(f)** The top 10 genes with the highest p-values among the 167 variable genes. Gene products were inferred from Marguerat et al ^5^.

The *S. pombe* transcriptome comprises a dataset of ∼7000 dimensions corresponding to expression levels (i.e., TPM) of ∼7000 annotated genes. By reducing the dimension from ∼7000 down to 2 in a stepwise manner, the data for all 64 samples could be plotted on a two-dimensional map, and the virtual timeline was drawn by monocle operations connecting proximal plots one by one (**Fig. 3b**). The plotted transcriptomes were divided into seven major clusters along the virtual timeline, with the first cluster (Cluster 1, **Fig. 3c**) corresponding to spores at the initial stage of germination. Cluster 1 was further divided into subclusters 1-a and 1-b (**Fig. 3c,d**), and we focused on genes for which expression varied significantly between these two subclusters (**Fig. 3e**). Statistical comparison of 167 genes revealed significant variation in expression (p < 0.05, **Fig. 3e**), and the top 10 most variable genes are listed in **Fig. 3f**.

#### Histone H3 fluctuates for germination

Within the gene list of **Fig. 3f**, we particularly focused on the gene *hht1* (encoding histone H3, copy 1; hereafter H3.c1); notably, *S. pombe* has three histone H3 genes (H3.c1, H3.c2, H3.c3), but it is unclear whether these genes are differentially used in *S. pombe*, as all three copies encode identical amino acid sequences ^11^. A monocle-based screen revealed that *hht1*/H3.c1 expression significantly varied upon germination, whereas the expression of *hht2*/H3.c2 nor *hht3*/H3.c3 did not change after germination. Variation of three H3 transcripts along the virtual timeline after refeeding demonstrated that the level of the H3.c1 mRNA decreased immediately after refeeding but later increased, probably reflecting the timing of DNA replication, whereas that of H3.c2 and H3.c3 remained essentially constant (**Fig. 4a-c**). These expression patterns are reminiscent of those previously reported for the H3.c1 and H3.c3 transcripts in proliferating cells, which showed a peak at S-phase, whereas H3.c2 stayed constant ^12^.

**Fig. 4.**
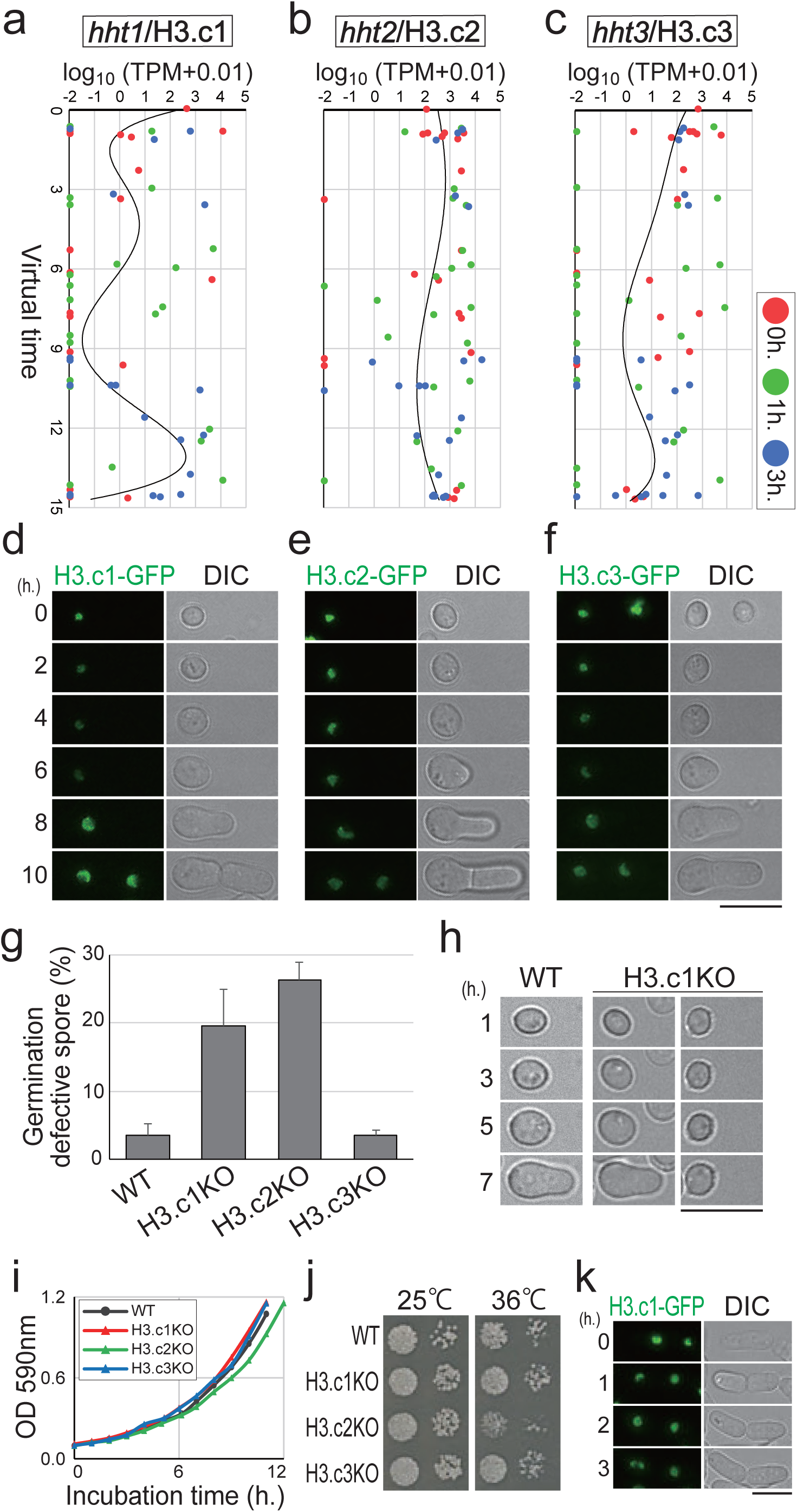
Expression of the histone H3 gene H3.c1 fluctuates during germination. The expression of one of the identified genes, *hht1* (H3.c1), indeed fluctuates. **(a-c)** *S. pombe* has three histone H3 genes: *hht1*/H3.c1 **(a)**, hht2/H3.c2 **(b)** and *hht3*/H3.c3 **(c)**. Expression of each histone H3 gene detected in 64 transcriptomes shown in Figure 3 was plotted versus the virtual timeline. Red, 0 h; green, 1 h; blue 3 h after feeding. **(d-f)** Each H3 protein fused with GFP was expressed under the native promoter at the endogenous level. Live-cell imaging started from dormant spores (0 h) until cells finished the first division. The fluorescence of only H3.c1-GFP fluctuated significantly (d). **(g,h)** Knock-out of H3.c1 (H3.c1KO) or H3.c2 causes defects in germination. **(g)** Frequencies of cells with defective germination in WT and knock-out strains of each H3 gene. Shown are live-cell differential interference-contrast (DIC) images of WT and H3.c1KO spores at germination. Hours (h.) indicate time after filming started. **(i, j)** H3.c1KO showed no visible growth defects during vegetative cycles, whereas H3.c2KO showed slight defects. H3.c2KO showed slight growth defects at 25°C **(i)**, which were even more evident on the plate at 36°C **(j). (k)** The level of H3.c1-GFP did not fluctuate during vegetative cycles in single-cell time-lapse observations. Scale bar, 10 µm.

To monitor protein levels of histone H3, a knock-in of the gene encoding GFP (green fluorescent protein) was carried out in *S. pombe*, resulting in expression of a *GFP* fusion with each of the genes encoding H3.c1, 2 and 3 under control of their native promoters (**Fig. 4d – f**). H3.c1-GFP fluorescence disappeared after germination was induced, but the fluorescence recovered to a considerable level at 8 h, when germ projection was evident (**Fig. 4d**). In contrast to the large fluctuations in the H3.c1-GFP signal, H3.c2-GFP fluorescence remained constant during germination, and H3.c3-GFP fluorescence fluctuated to a lesser extent than did that of H3.c1-GFP (**Fig. 4e,f**). Indeed, time-lapse behaviour of all H3 proteins was consistent with the kinetics predicted for each of the H3 mRNAs (**Fig. 4a-c**), validating our estimation of the virtual time-line.

We then tested whether the three H3 proteins contributed to spore germination. Both H3.c1 knock-out (H3.c1KO) and H3.c2KO spores exhibited defective germination, as indicated by a significant delay or stall in the formation of the germ projection, whereas H3.c3KO spores germinated almost normally (**Fig. 4g,h**). In contrast to the germination defects, H3.c1KO cells did not exhibit visible growth defects during mitotic cell cycle, whereas H3.c2KO showed temperature sensitivity and slight growth defects during the mitotic cycle (**Fig. 4i,j**) ^12^. In line with these results, the fluctuation in H3.c1-GFP fluorescence seen during germination was not evident in the mitotic cycle (**Fig. 4k**), demonstrating that H3.c1 expression is modulated specifically during germination. We thus postulated that H3.c1 is not just a redundant histone H3 but rather a specific variant that plays a role in promoting germination, whereas H3.c2 is a general and housekeeping histone expressed and required at all time.

Because the amino acid sequences of the three H3.c1–3 proteins are identical, we speculated that the observed differential expression of H3.c1 and H3.c2 was a consequence of differences in their DNA sequences in the 5’ promoter or coding region. The 3’ terminator region cannot account for the difference because each of the engineered fusion genes (H3.c1-GFP, etc.) contained the same exogenous terminator (T_ADH_), and indeed their expression patterns were unaffected by this (see **Fig. 4d**). We therefore replaced the promoter for H3.c1 (P_H3.c1_) with the one for H3.c2 (P_H3.c2_) at the endogenous locus (the strain “P_H3.c1_ → P_H3.c2_”, **Fig. 5a**) to evaluate the effect of P_H3.c1_ on the kinetics of H3.c1 expression. After inducing germination in WT cells, the level of H3.c1-GFP gradually decreased but then increased significantly after 5 h (**Figs. 4d–f** and **5a,b**). In contrast, this increase was not observed when P_H3.c1_ was replaced with P_H3.c2_ (P_H3.c1_ → P_H3.c2_, **Fig. 5a,b**), indicating that the promoter region of H3.c1 is responsible for the observed transcriptional boost of H3.c1 during germination.

**Fig. 5.**
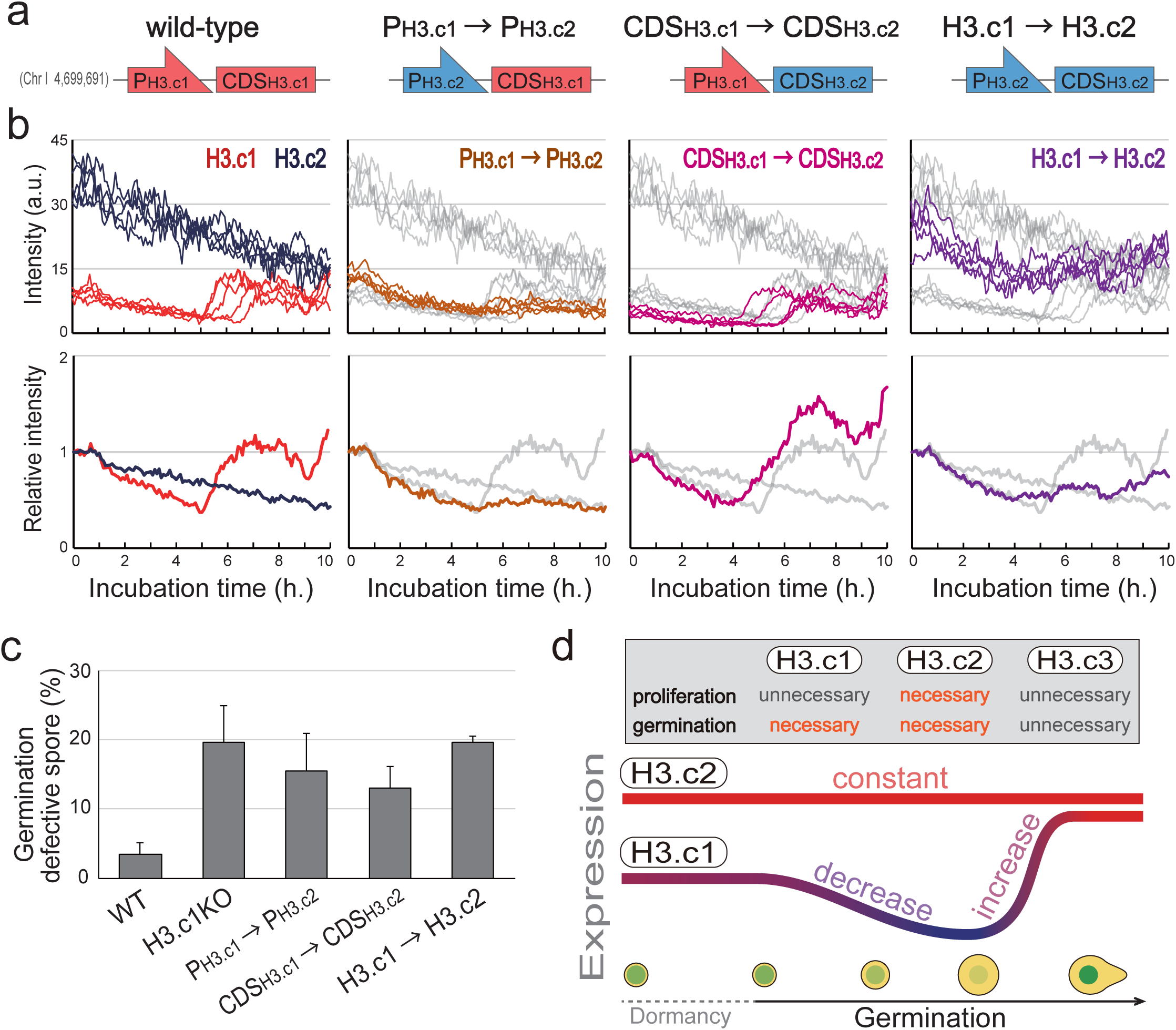
Control of H3.c1 expression by its promoter is key for fluctuation. Temporal changes in the levels of the various histone H3 proteins are controlled by their promoters (P) and coding sequences (CDS). **(a)** Schematics illustrating states of H3.c1 and H3.c2 gene loci used in this study. For example, P_H3.c1_ → P_H3.c2_: the strain in which the promoter of H3.c1 (P_H3.c1_) was replaced with that of H3.c2 (P_H3.c2_); H3.c1 → H3.c2: both P and CDS were exchanged. **(b)** Fluorescence intensity of GFP was measured over time. Top, absolute values are shown; bottom, intensities were normalised to values at time 0 (start of observation). Intensities of WT H3.c1-GFP (red) and H3.c2-GFP (dark blue) are shown to the left, which are also copied onto other graphs (grey traces). Similarly, P_H3.c1_ → P_H3.c2_-GFP (brown), CDS_H3.c1_ → CDS_H3.c2_-GFP (yellow), and H3.c1 → H3.c2-GFP (magenta) are shown. **(c)** Artificial alteration of H3.c1 expression caused defects in germination. Error bars, s.d. **(d)** A model for histone H3.c1 expression for efficient germination. Expression decreases once in the initial stage but is substantially boosted in the later stages of germination.

Strain P_H3.c1_ → P_H3.c2_ displayed defects in spore germination, indicating that the observed transcriptional boost ensured that germination would progress (**Fig. 5c**). Replacement of the coding sequence (CDS) of H3.c1 with that of H3.c2 (“CDS_H3.c1_ → CDS_H3.c2_”, **Fig. 5a**) reduced the expression even at the very initial stage of germination, which might account for the minor defects in germination observed for this mutant, as the boost at later stages occurred normally (**Fig. 5b,c**). Replacement of both the promoter and coding regions of H3.c1 by those of H3.c2 (“H3.c1 → H3.c2”, **Fig. 5a**) resulted in its constitutive expression, which was comparable with the expression kinetics of endogenous H3.c2 (**Fig. 5b**). This implied that the total H3 level in this strain was consistently high even from the initiation stage of germination. This strain had a high rate of defective germination (**Fig. 5c**), demonstrating that the cells needed to repress the H3 expression to avoid an overabundance early during germination, whereas there was need to boost expression at a later stage around DNA replication. Thus, spores emerge from dormancy via temporal upregulation of a specific H3 gene using the promoter and coding regions of that H3 gene, even though all three H3 proteins have identical amino acid sequences (**Fig. 5d**).

#### Power of time-lapse, single-cell profiling

We took the approach of single cell–based transcriptomes to investigate the long-lasting issue of how the dormancy of spores is broken in response to environmental change. Only a single example of *S. pombe* single-cell transcriptomics has been published ^4^; by comparison, our method entails easier handling procedures and requires no specialised devices, and indeed it yields a higher rate of gene detection than the previous protocol. Statistical comparison of gene-expression profiles created from a single cell and from bulk cells demonstrated that the validity of the single-cell protocol matches that of bulk cells (see **Fig. 1e,f**). As a proof of principle, we utilised bioinformatics to align transcriptomes from single cells at various stages of germination along a virtual timeline. This analysis highlighted *hht1*, one of three histone H3 genes in *S. pombe*, as a candidate gene for which expression is altered at the initial stage of germination (see **Fig. 3**). Analysis of the temporal kinetics of mRNA levels estimated along the virtual timeline revealed a correlation with the fluorescence emitted by histone H3-GFP proteins as assessed with live-cell imaging, further validating our methodology (**Fig. 4a–f**). Based on expression data acquired over time (**Fig. 3c**), the 64 transcriptomes we created from single cells comprised 7 clusters that reflected the various stages of germination. Indeed, each stage may reflect the progress of stepwise cellular events, e.g., morphological alterations and DNA replication prior to entry into proliferation cycles.

#### Histone H3 drives cell-cycle initiation

Our study demonstrates functional differences between the three copies of histone H3 genes encoding identical amino acid sequences. Hht1/H3.c1 and Hht2/H3.c2 play major roles in germination, whereas Hht3/H3.c3 is dispensable. Previous studies revealed that expression of both H3.c1 and H3.c2 in proliferating cells peaked in S phase ^12^. We, however, found that the H3.c1 level decreased in the initial stage of germination and then was upregulated in later stages, whereas H3.c2 was continuously expressed at a high level (**Fig. 5a,b**), indicating that their expression during germination is regulated in a manner distinct from that of mitotic cycles. In previous transcriptome analyses using cells that were shifted from G0 quiescence to the cycling stage, no H3 genes were found to have altered transcription ^13^, suggesting that changes in H3.c1 expression is specific in spore germination. We envision that at the end of gametogenesis, spores fix their chromatin a state to ensure survival during prolonged dormancy, and this state must be reset upon germination via the downregulation of H3.c1 expression, which allows yeast cells to establish a nutrient-replete cellular programme and proliferate.

We propose that cooperative expression of H3.c2 (continuous expression) and H3.c1 (variable) is essential for efficient germination (**Fig. 5d**). In this regard, within dormant spores, H3.c1 and H3.c2 are abundant in nucleosomes, but H3.c1 must be downregulated at the initial stage of germination in response to changes in nutrient availability, which may contribute to a relaxation of chromatin structure to enable a global transcriptional boost. Meanwhile, H3.c2 remains abundant, possibly serving as a pool of housekeeping histones for nucleosome assembly.

As germination proceeds, H3.c1 level must be increased to associate with newly synthesised DNA during S phase of the pioneer round of the cell cycle. The transcriptional boost can be ascribed to the promoter activity of the H3.c1 gene (**Fig. 5a,b**). In contrast, the H3.c1 CDS appears to be required for maintenance of its expression to an optimal level from the very initial stage of germination, possibly because transcripts that contain the H3.c1 CDS might be more stable than those that contain the H3.c2 CDS at that time. Alternatively, the chromatin region that includes the H3.c1 locus may be silenced (e.g., facultative heterochromatin), and the H3.c1 CDS might open the chromatin to facilitate transcription of the H3.c1 gene at the later stage of germination.

At early points in the development of mammalian embryos, the genome-wide chromatin state (i.e., open or closed) may change dramatically to promote cell division ^14,15^. The fission yeast is a simple and unique organism with respect to H3 structure, as all three copies are composed of identical amino acid sequences that are a hybrid of the canonical mammalian H3.2 and H3.3 sequences, which is unlike higher eukaryotes that have three distinct H3 sequences ^16^. Hence, the evolutionarily simple fission yeast uses its three H3s in different ways by modulating the temporal kinetics of transcription of their individual genes to adapt to environmental changes and enter a routine mitotic state.

## Acknowledgements

We thank Taro Nakamura and Chikashi Shimoda for advice on fission yeast spore biology, Sachiyo Aburatani and Hisashi Anbutsu for support in OIL, Toru Maruyama for advice on informatics, and Kiyofumi Takahashi and Chikako Sakanashi for technical support in RNA-seq. This study was supported by JSPS KAKENHI JP25291041, JP15H01359, JP16H04787, JP16H01317 and JP18K19347 to M.S. This study was also supported by The Uehara Memorial Foundation and by Waseda University grants for Special Research Projects 2017B-242, 2017B-243 and 2018B-222 to M.S. This work was partially supported by the JSPS Core-to-Core Program, A. Advanced Research Networks, and by the Platform Project for Supporting Drug Discovery and Life Science Research (Basis for Supporting Innovative Drug Discovery and Life Science Research (BINDS)) from Waseda University under Grant Number JP19am0101104.

## Author contributions

H.Ts. conducted experiments using fission yeast supervised by M.S. and performed bioinformatic operations supervised by K.A. M.H., T.Y., K.A. and H.Ta. designed the single-cell transcriptome methodologies and performed RNA-seq studies. N.O. provided initial guidance for the project, which was conceived and designed by M.S.

## Methods online

### Yeast strains, media and genetics

Standard materials and methods were used for *S. pombe* biology ^17^. Strains used in this study are listed in **Supplementary Table 1** online. Briefly, YE5S (yeast extract with supplements) was used for vegetative growth and also as refeeding medium to induce the germination of spores. Edinburgh minimal medium supplemented with nitrogen sources and amino acids was used for the preparation of vegetative cells for RNA sequencing. Sporulation agar plates were used for induction of mating and meiosis.

The prototrophic strain (L975: h90 WT) was mainly used as a host strain for genome editing and also as a control strain for assessing germination, because background mutations such as those for leucine or uracil autotrophy can sometimes affect germination.

Conventional methods were used for gene knock-out and knock-in ^18-20^. In GFP-tagged strains, *GFP* was knocked in at the 3’-end of the gene of interest. For instance, in strain hht1(H3.c1)-GFP, *GFP* was inserted at the 3’-end of the coding sequence of *hht1* (in frame) to replace the endogenous *hht1* with the *hht1*-GFP fusion gene. This ensured that the expression of *hht1-GFP* remained under control of the native *hht1* promoter. Strain Pnmt41-*bgs2* was created using the template plasmid pFA6a-bsdMX6-pnmt41, in which the kanMX6 of pFA6a-kanMX6-pnmt41 was replaced with bsdMX6. The Pnmt41 promoter is a thiamine-repressible promoter, which moderately induces expression of the downstream gene in medium lacking thiamine ^21^.

For vegetative growth assays, cells were grown in YE5S liquid medium at 25°C overnight until 10^6^–10^7^ cells/ml was reached (“precultures”). For liquid culture–based assays, the preculture was inoculated into 10 ml YE5S medium, which was then placed in a shaker incubator (130 rpm) at 25°C overnight. OD_590_ values were measured every hour using a spectrophotometer (GeneQuant 1300; GE Healthcare Life Sciences). For plate-based growth assays, the preculture was concentrated through mild centrifugation to 2 × 10^8^cells/ml, after which 10-fold serial dilutions were made. Approximately 10^2^–10^3^cells were spotted onto YE5S plates followed by incubation at 25°C or 36°C.

### Preparing spheroplasts

Fission yeast cells cultured in appropriate media were collected and washed once with Buffer A (1.2 M D-sorbitol, 50 mM citrate-phosphate, pH 5.6). Here we used cells expressing Ste11-GFP to pick up non-starved cells ^22^. Cells were then suspended in 200 *µ*l Buffer A containing 200 mg lysing enzyme from *Trichoderma harzianum* (Novozymes) and incubated at 32°C for 30 min with stirring every 10 min. Cells were then washed three times with Buffer B (1.2 M D-sorbitol, 30 mM Tris-HCl, pH 7.6) and finally suspended with 50 *µ*l Buffer B.

### Preparing spores, and induction of germination

Pnmt41-*bgs2* mutant cells (for main experiments) as well as WT (expressing Ste11-GFP) were used for preparation of spores. Briefly, sexual differentiation (mating, meiosis and sporulation) was induced on solid sporulation agar plates without nitrogen sources for 3 days at 30°C until spores were spontaneously separated. For induction of spore germination, cells containing spores were washed once with Buffer B and suspended into rich medium (i.e., YE5S) for 0–3 h (feeding) at 30°C. To pick up single spores, cells at each time point were collected and washed with Buffer B and then resuspended into 500 *µ*l Buffer B prior to pick-up.

### Single-cell pick-up

Methods for the entire procedure are as follows. First, a bulk culture of cells was prepared in Edinburgh minimal medium (liquid) or on sporulation agar plates depending on experimental needs. Cells were collected and spheroplasts produced if necessary. Each cell suspension prepared in the previous step was mounted on a 35-mm dish and placed on the stage of a light microscope. With visual inspection through the eyepiece lenses, a Drummond Microdispenser (Drummond Scientific Company) pipette was used so that a single cell could be absorbed with 1 *µ*l of buffer. Each single cell was then transferred into a 0.2-ml tube containing 2.5 *µ*l Collection Solution (2 U/*µ*l RNase inhibitor and 0.2% (w/v) NP-40 in RNase-free ddH_2_O). Thus, each sample was expected to contain a single cell in 3.5 *µ*l liquid in total. Once collection was done, samples were stored at –80°C.

### RNA extraction and cDNA preparation

For preparation of sequence libraries, samples were used for cDNA preparation with the “bead-seq” method ^6^. Single-cell samples were defrosted, and 6.5 *µ*l lysis solution was added; the final concentrations of reagents were as follows: 0.45 × PCR buffer II (Life Technologies), 0.45% (w/v) NP-40, 4.5 mM dithiothreitol, 0.18 U/*µ*l SUPERase•In (Ambion), 0.36 U/*µ*l RNase inhibitor (Ambion), 0.15mM each dNTP, dual-biotinylated anchored “oligo 1” primers immobilised on ∼10^7^ beads/sample ^6^. The sequences of the oligonucleotides used are listed in the **Supplementary Table 2**. Samples were transferred into a Caliper Zephyr Compact Liquid Handler (Perkin Elmer, CA, USA) coupled with a homemade operation program based on the Bead-seq method. Samples were then incubated by 70°C for 5 min and chilled to RT, after which 5 *µ*l reverse transcription solution was added (final concentration: 0.45 × PCR buffer II, 1.35 mM MgCl_2_, 8 U/*µ*l Superscript III reverse transcriptase from Life Technologies, and 0.4 U/*µ*l RNase inhibitor). Samples were incubated at 55°C for 30 min followed by 70°C for 5 min. Beads were magnetically-collected in tube and washed with T&T buffer (0.1% (w/v) Tween 20, 10 mM Tris-HCl pH 8.0) and resuspended in 12 *µ*l T&T buffer.

After reverse transcription, 6 *µ*l of tailing solution (0.5× PCR buffer II, 0.75 mM MgCl_2_, 1.5 mM dATP, 0.15 U/*µ*l RNase H, 0.188 U/*µ*l terminal deoxynucleotidyl transferase) was added to the samples, with subsequent incubation in a shaker at 37°C for 15 min followed by 70°C for 10 min. Samples were then washed with T&T buffer once and resuspended with 12 *µ*l T&T buffer. For synthesis of the second strand, 19 *µ*l of each reaction mixture (final concentration: 1× Ex Taq buffer (TaKaRa Bio), 0.12 mM dNTP, 0.49 *µ*M anchored “oligo 2” primers, 0.05 U/*µ*l Ex Taq Hot Start version (TaKaRa Bio)) was added, and the samples were applied to a thermal cycler: 95°C for 30 s, 44°C for 5 min, and 72°C for 6 min, followed by cooling to 4°C.

Synthesised cDNA was then amplified via two-step PCR. For the first step, 1 *µ*l of amplification solution 1 (1× Ex Taq buffer, 0.25 mM dNTP, 0.05 U/*µ*l Ex Taq Hot Start version, 0.49 *µ*M anchored “oligo1_T15” primer) was added to each sample, and thermal cycling was performed as follows: boiling at 95°C for 0.5 min, followed by 18 cycles of 95°C for 0.5 min, 67°C for 1 min and 72°C for 6 min, followed by cooling to 4°C. Samples were then washed twice with 0.6 × AMPure XP reagent (Beckman Coulter) and resuspended with 20 *µ*l T&T buffer.

For the second PCR step, 5 *µ*l of each sample from the first PCR step was mixed with 45 *µ*l of amplification solution 2 (1× Ex Taq buffer, 0.25 mM dNTP, 0.05 U/*µ*l Ex Taq Hot Start version, 1 *µ*M anchored “oligo1_T15” primer, 1 *µ*M anchored “oligo 2” primers), and thermal cycling was performed as follows: boiling at 95°C for 0.5 min, and 12 cycles of 95°C for 0.5 min, 67°C for 1 min and 72°C for 6 min, followed by cooling to 4°C. Samples were then washed once with 0.6 × AMPure XP reagent (Beckman Coulter) and resuspended with 20 *µ*l T&T buffer.

### Sequencing

Amplified cDNA products were then fragmented for next-generation sequencing using the Nextera XT DNA Sample Prep kit (illumina). The resultant sequencing pool was subjected to TapeStation 4200 electrophoresis using High Sensitivity D5000 SceenTape (Agilent technology). Next-generation sequencing was then performed with HiSeq 2500 (illumina; performed by Macrogen Corp., Japan). Pair-end reads of 101 nucleotides were sequenced.

### Computational analysis for gene expression levels

Read-out sequences in FASTQ-format files were mapped onto the reference genome of *S. pombe*: Schizosaccharomyces_pombe.ASM294v2.25. Read counts were calculated with RSEM-1.3.0 ^23^ and normalised to scores of transcripts per million mapped reads (TPM) ^24^.

### Estimation for the virtual time-line

For the drawing of scatter plots (**Figs. 1e,f** and **2c**), the software Subio Platform ver.1.19 (Subio Inc., Amami, Japan) was used.

For virtual time-line ordering analysis (**Fig. 3b**), we utilised the R package monocle ver.2.4.0 ^8-10^. Matrices for TPM, for cell attributes, and for gene annotation were used as input data; a TPM matrix was defined as a matrix for TPM values, in which each row corresponds to each gene and each column to each single cell. A cell attribute matrix was defined as values for experimental conditions such as time after feeding (time for germination induction). Each row corresponds to a single cell and each column to a particular condition. A gene annotation matrix was defined as a gene information dataset, in which each row corresponds to a single gene and each column to annotated information. Input data were used for dimensionality reduction and virtual timeline deduction with reverse graph embedding methods ^10^. For the monocle operation, standard parameters were used according to the instructions of the supplier, and genes with little or no expression (below 0.1 RNA copies per cell (RPC) and genes detected in fewer than 10 samples were excluded from the operation.

For clustering of transcriptome datasets (**Fig. 3c**), the R package mclust ver.5.3^25,26^ was used. The output dataset of the aforementioned monocle operation was used for mclust input. Samples were classified through Gaussian kernel density estimation. For further clustering using dendrograms (**Fig. 3d**), the R package amap ver.0.8.14 (https://CRAN.R-project.org/package=amap), and dendextend ver. 1.8.0 ^27^ were used. The TPM matrices belonging to transcriptomes in Cluster 1, as classified by mclust, were used for input.

Similarity between samples (**Figs. 2d** and **3d**) was estimated by measuring the Euclidean distance followed by hierarchical clustering with Ward’s method. Based on the similarity, two transcriptomes with the closest gene expression profiles were connected by a branch, the size of which is denoted as ‘height’. A pair of samples with a relatively low height value could be interpreted as having greater similarity.

For differential expression analysis (**Fig. 3e**), the R package edgeR ver. 3.18.1 ^28,29^ was used. The read count matrices belonging to Cluster 1, as classified by mclust, were used for input. A read count matrix was defined as a dataset for read count for next-generation sequencing, in which each row corresponds to a single gene and each column to a single cell. Samples in Cluster 1 were divided into two subclusters, 1-a and 1-b. The difference between the expression level of a particular gene in each of these subclusters was statistically examined with an exact test. logFC = log (average transcript level in subcluster 1-b) – log (average transcript level in subcluster 1-a). logCPM = {log (average transcript level in subcluster 1-a) + log (average transcript level in subcluster 1-b)} / 2. FC, fold change; CPM, counts per million mapped reads.

### Microscopy

For live-cell imaging of green fluorescent protein (GFP) and differential interference-contrast microscopy, we applied our published standard methods ^20^. Briefly, the observation system comprised the DeltaVision-SoftWoRx image acquisition system (Applied Precision) equipped with the Olympus inverted microscopes IX71 and IX81 and CoolSNAP HQ2 CCD cameras (Photometrics). In **Fig. 4d–f**, the parental strain was induced for sporulation for 3 days at 26.5°C, and spores were suspended with 200 *µ*l ddH_2_O and mounted onto a glass-bottomed dish (Iwaki glass) precoated with lectin from *Glycine max* (Sigma). The dish was filled with 2 ml YE5S prior to observation. To measure GFP fluorescence, images for 20 sections along the Z axis (0.2 *µ*m interval) were filmed, deconvolved, and finally projected to a single plane using SoftWoRx software.

## Supplementary information

**Supplementary Table 1.**
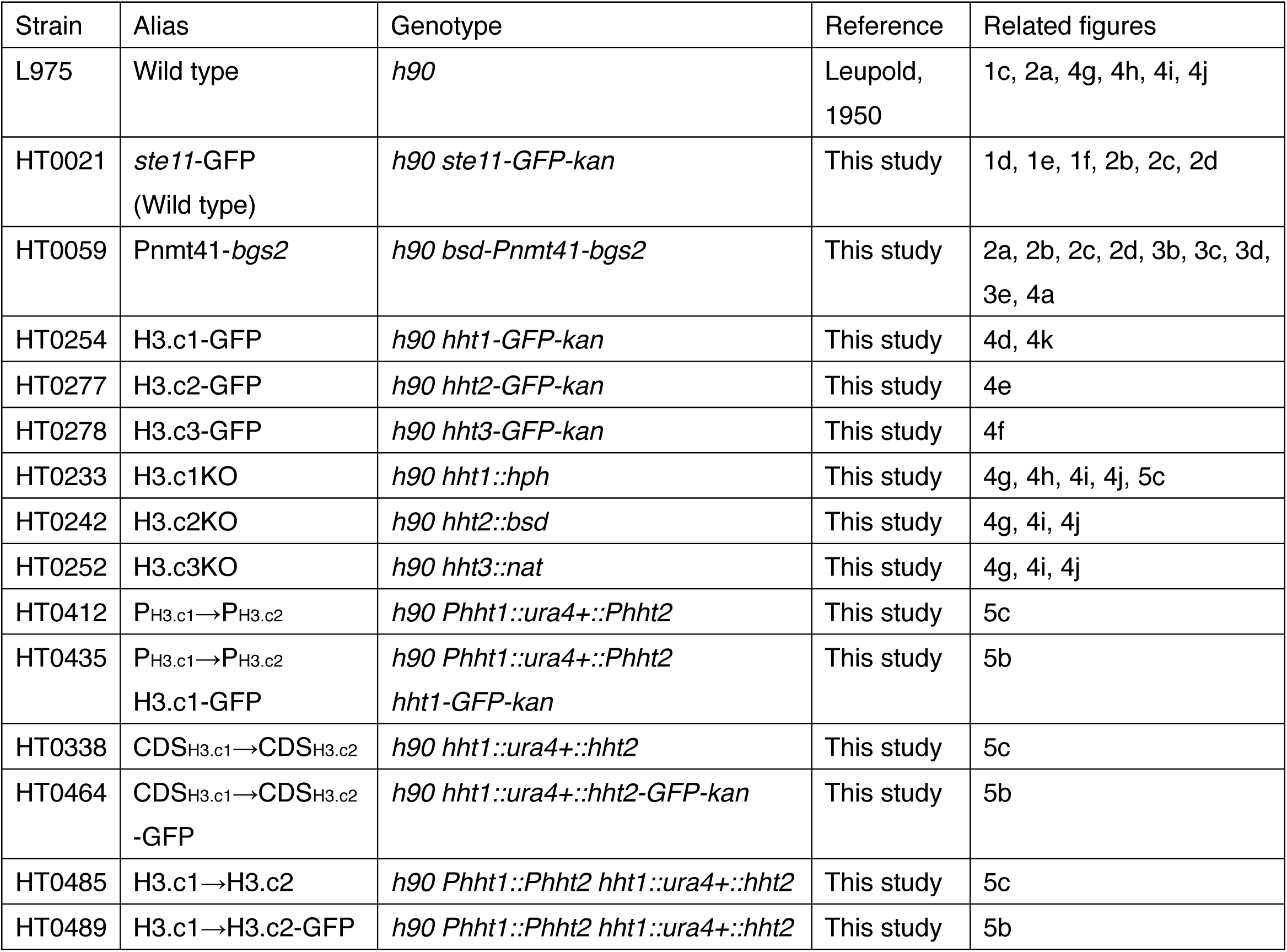
Strains used in this study.

| Strain | Alias | Genotype | Reference | Related figures |
| --- | --- | --- | --- | --- |
| L975 | Wild type | *h90* | Leupold, 1950 | 1c, 2a, 4g, 4h, 4i, 4j |
| HT0021 | *ste11*-GFP<br>(Wild type) | *h90 ste11-GFP-kan* | This study | 1d, 1e, 1f, 2b, 2c, 2d |
| HT0059 | *Pnmt41-bgs2* | *h90 bsd-Pnmt41-bgs2* | This study | 2a, 2b, 2c, 2d, 3b, 3c, 3d, 3e, 4a |
| HT0254 | H3.c1-GFP | *h90 hht1-GFP-kan* | This study | 4d, 4k |
| HT0277 | H3.c2-GFP | *h90 hht2-GFP-kan* | This study | 4e |
| HT0278 | H3.c3-GFP | *h90 hht3-GFP-kan* | This study | 4f |
| HT0233 | H3.c1KO | *h90 hht1::hph* | This study | 4g, 4h, 4i, 4j, 5c |
| HT0242 | H3.c2KO | *h90 hht2::bsd* | This study | 4g, 4i, 4j |
| HT0252 | H3.c3KO | *h90 hht3::nat* | This study | 4g, 4i, 4j |
| HT0412 | P<sub>H3.c1</sub>→P<sub>H3.c2</sub> | *h90 Phht1::ura4+::Phht2* | This study | 5c |
| HT0435 | P<sub>H3.c1</sub>→P<sub>H3.c2</sub><br>H3.c1-GFP | *h90 Phht1::ura4+::Phht2*<br>*hht1-GFP-kan* | This study | 5b |
| HT0338 | CDS<sub>H3.c1</sub>→CDS<sub>H3.c2</sub> | *h90 hht1::ura4+::hht2* | This study | 5c |
| HT0464 | CDS<sub>H3.c1</sub>→CDS<sub>H3.c2</sub><br>-GFP | *h90 hht1::ura4+::hht2-GFP-kan* | This study | 5b |
| HT0485 | H3.c1→H3.c2 | *h90 Phht1::Phht2 hht1::ura4+::hht2* | This study | 5c |
| HT0489 | H3.c1→H3.c2-GFP | *h90 Phht1::Phht2 hht1::ura4+::hht2* | This study | 5b |

**Supplementary Table 2.**
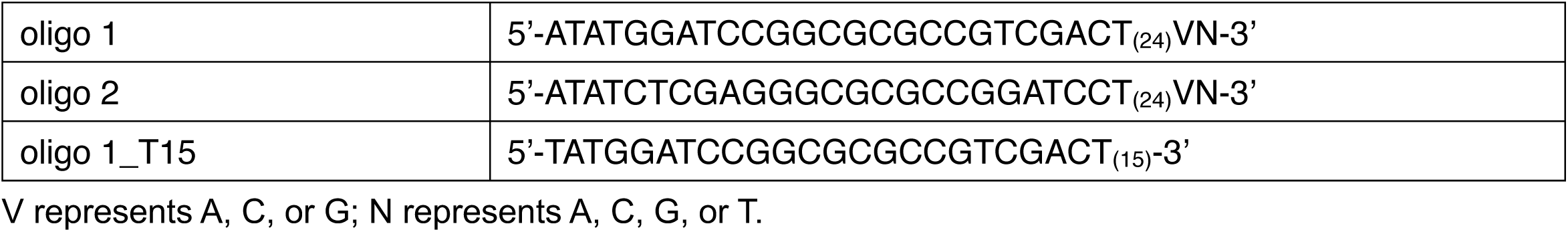
Oligonucleotides used in this study.

**Supplementary Fig. 1.**
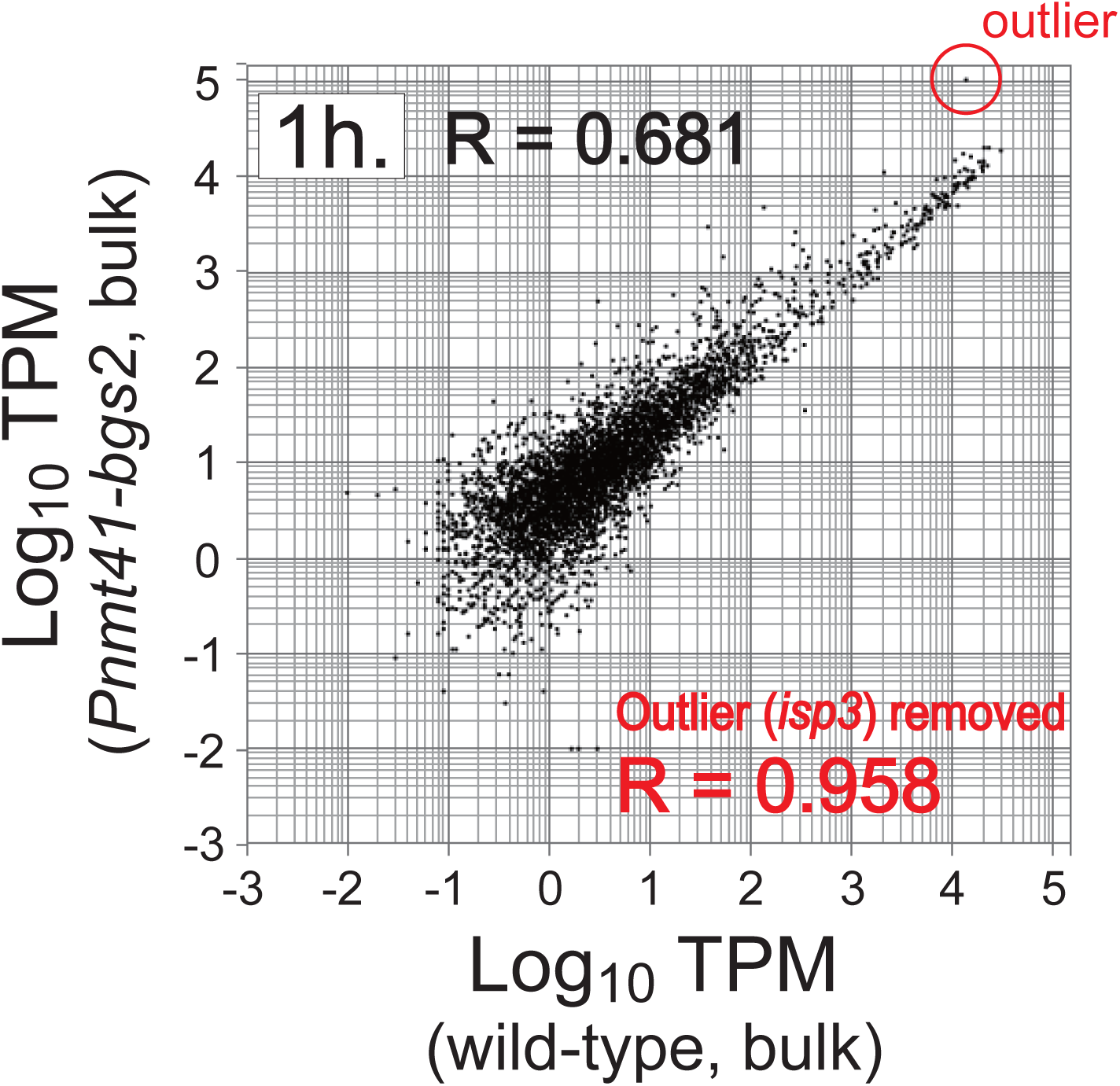
Comparison of transcriptomes derived from cultures of bulk spores. This figure provides supplementary information for **Fig. 2d**, in which the correlation coefficient (R value) was 0.681 between transcriptomes from bulk WT and Pnmt41-*bgs2* cells. The red circle denotes the single outlier gene, *isp3*. The similarity between these two profiles increased (R = 0.958) when the data for *isp3* was not considered. We thus concluded that reduction of cellular Bgs2 in Pnmt41-*bgs2* cells did not affect the overall transcription profile except for that of *isp3*. Because Isp3 is a component of the surface layer of the spore wall (Fukunishi et al., Mol. Biol. Cell., 2014), a reduced level of Bgs2 may alter spore-wall integrity.

